# Image Correlation Spectroscopy is a Robust Tool to Quantify Cellular DNA Damage Response

**DOI:** 10.1101/2024.08.05.606697

**Authors:** Angelica A. Gopal, Bianca Fernandez, Paul W. Wiseman, J. Matthew Dubach

## Abstract

The DNA Damage response (DDR) is both essential and highly complex. Evaluating the DDR is a critical aspect of cell biology. Counting DNA damage foci is one of the most common approaches to study the DDR. Yet, quantification of protein foci suffers from experimental limitations, subjectivity of analysis and is restricted to a handful of the hundreds of DDR proteins. Here we apply image correlation spectroscopy (ICS) to quantify the local clustering at sites of DNA damage directly. We found that ICS outperformed foci counting of traditional DDR markers and enabled quantification of other markers without the complex labeling procedures that are otherwise required. ICS analysis also provided insight into DDR protein recruitment that was previously undetectable. Further expansion incorporating analysis to cell cycle classification demonstrates a rapid, non-biased approach to fully study the DNA damage response within cells. ICS analysis presents an objective, quantitative image analysis technique to study the DNA damage response in unaltered cells that we expect will significantly enhance quantitative DNA damage response research.

## Introduction

The DNA damage response (DDR) represents multiple different pathways that are dependent on several factors including cell cycle and the type of damage^1-3^. DNA damage and the DDR are prominent in several disease areas such as cancer^4^, neurodegeneration^5^ and aging^6^, and the aberrant DDR is a major target of cancer therapies^7^. Collectively, the DDR comprises hundreds of proteins that are recruited to sites of damage in a concerted manner serving as scaffolds, DNA binders, modifying enzymes and repair enzymes. Largely due to the massive evolutionary pressure to maintain the genome, the DDR is enormously complex, rendering it difficult to study in cells.

Current methods to evaluate DNA damage vary from molecular strategies such as polymerase chain reaction^8^, comet assays^9^ and gel electrophoresis^10^ to intracellular approaches that include, fluorescence in situ hybridization^11^, radioimmunoassay^12^ and TUNEL assays^13^. Analytical methods have also been employed, for example, gas or liquid chromatography, mass spectrometry and electrochemical approaches have been used to identify DDR components and signaling^7^. However, the gold standard for assessing DNA damage levels in cells is fluorescence imaging and counting DNA repair foci formation by labeling established protein markers, such as Ku70/80^14^, γH2AX^15^ or XRCC1^16^, through immunofluorescence or genetically encoded fluorescent proteins.

Historically, foci counting was performed manually and relied on individual researchers to identify foci, introducing an inherent bias to the process. More recently, foci counting has been widely automated via computer-based algorithms^17,18^. Numerous software tools have been released to count foci via methods of signal thresholding, maxima finding, and morphological reconstructions to accurately detect foci in nuclei. However, foci are not often individually distinguishable and are present at varying sizes and intensities, leading to some ambiguity in thresholding, segmenting and counting parameters, with a wide variety of software employing different algorithms^19-21^. Furthermore, γH2AX foci were recently revealed to represent clustered structures of γH2AX nano-domains^22^, suggesting multiple sites of DNA damage are present in single foci. Thus, it is likely that not all foci are the same – larger foci representing a greater degree of or more mature DNA damage than smaller foci.

In addition to measurements of known foci markers, studies using laser micro-irradiation have shown that hundreds of different proteins are recruited to sites of DNA damage^23^, despite not forming foci that are resolvable in fluorescence images when more biologically relevant DNA damage is induced. Therefore, the role of the majority of DDR proteins remains poorly understood because their recruitment to DNA damage foci is difficult to quantify with traditional approaches. Typically, non-recruited protein is pre-extracted, through lysis of the cell and nucleus, prior to fixation to reduce the background concentration and enhance the resolution of protein foci. However, the demonstration that DNA damage foci are biomolecular condensates where foci protein concentration exists in equilibrium with the nuclear concentration^24^ suggests that pre-extraction may impact the DNA damage foci as well to hinder the accuracy of the results.

To overcome the limitations of current cellular DDR analysis approaches, we hypothesized that measuring clustering of labeled proteins within the nucleus could quantify the degree of DNA damage and involvement of any DDR protein. To test this approach, we employed fluorescence microscopy imaging with fluorescence fluctuation analysis via spatial image correlation spectroscopy (ICS) as a tool for measuring the clustering of proteins during DDR in multiple cancer cell lines treated with DNA damaging agents and DDR therapeutics. Fluorescence correlation spectroscopy (FCS), which requires temporal imaging, has been previously used for the measurements of macromolecular aggregation via autocorrelation analysis of fluorescence intensity fluctuations that arise due to changes in the local concentration of fluorescence molecules in a tiny focal volume^25-27^. Thus, FCS is well suited to study single, well-defined foci. Conversely, ICS was developed as the imaging analog of FCS to measure densities and aggregation states of membrane receptors from spatial correlation analysis of very small fluorescence fluctuations in single images^28,29^. Thus, ICS presents a potential alternative method to quantify DNA damage and expand the detection of DDR response elements beyond established protein markers.

## Methods

### Theory

Image correlation spectroscopy is based on the analysis of fluorescence intensity fluctuations arising from variations in the number of fluorescent particles within focal spots imaged in space using a fluorescence microscope^30^. Under the assumptions of particle ideality and linear fluorescence emission (i.e., no signal saturation or energy transfer), the detected mean fluorescence intensity of tagged molecules varies linearly with the concentration of fluorophores in a focal spot (Fig. 2A-B). The square relative intensity fluctuation (intensity variance/mean intensity, Eq. 1) will be the mean number of detected independent fluorescent particles per focal spot, since ideal behavior entails that the molecules obey Poisson statistics within the volume:

**Figure 1.**
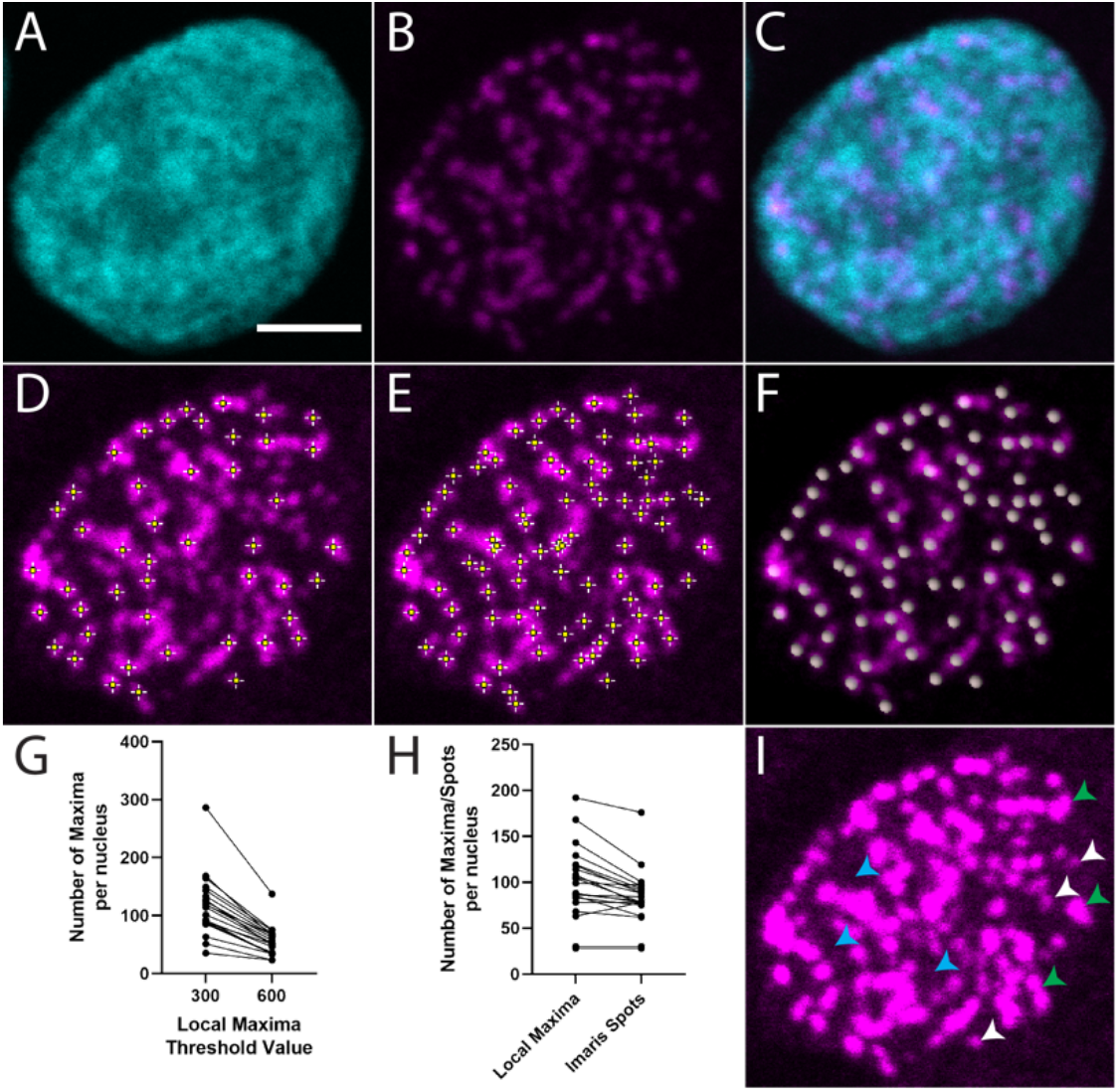
Common approaches to count DNA damage foci. (**A**) DAPI channel with 5 µm scale bar, (**B**) γH2AX channel, (**C**) Composite of A) and B). (**D-E**) Foci counting with Find Maxima tool in ImageJ with a threshold of D) 600 and E) 300. (**F**) Foci counting using Imaris Spots with an automatic threshold and an expected spot size of 1 µm. (**G**) Number of maxima per nucleus from different thresholds using ImageJ for the same cell. (**H**) Foci counting comparison between a user defined threshold in ImageJ and automatic threshold in Imaris on the same cell. (**I**) Intensity adjusted γH2AX channel showing lower intensity foci (white arrows), larger foci (green arrows) and higher background intensities (blue arrows). Figure S1 provides larger images for further clarity.

**Figure 2.**
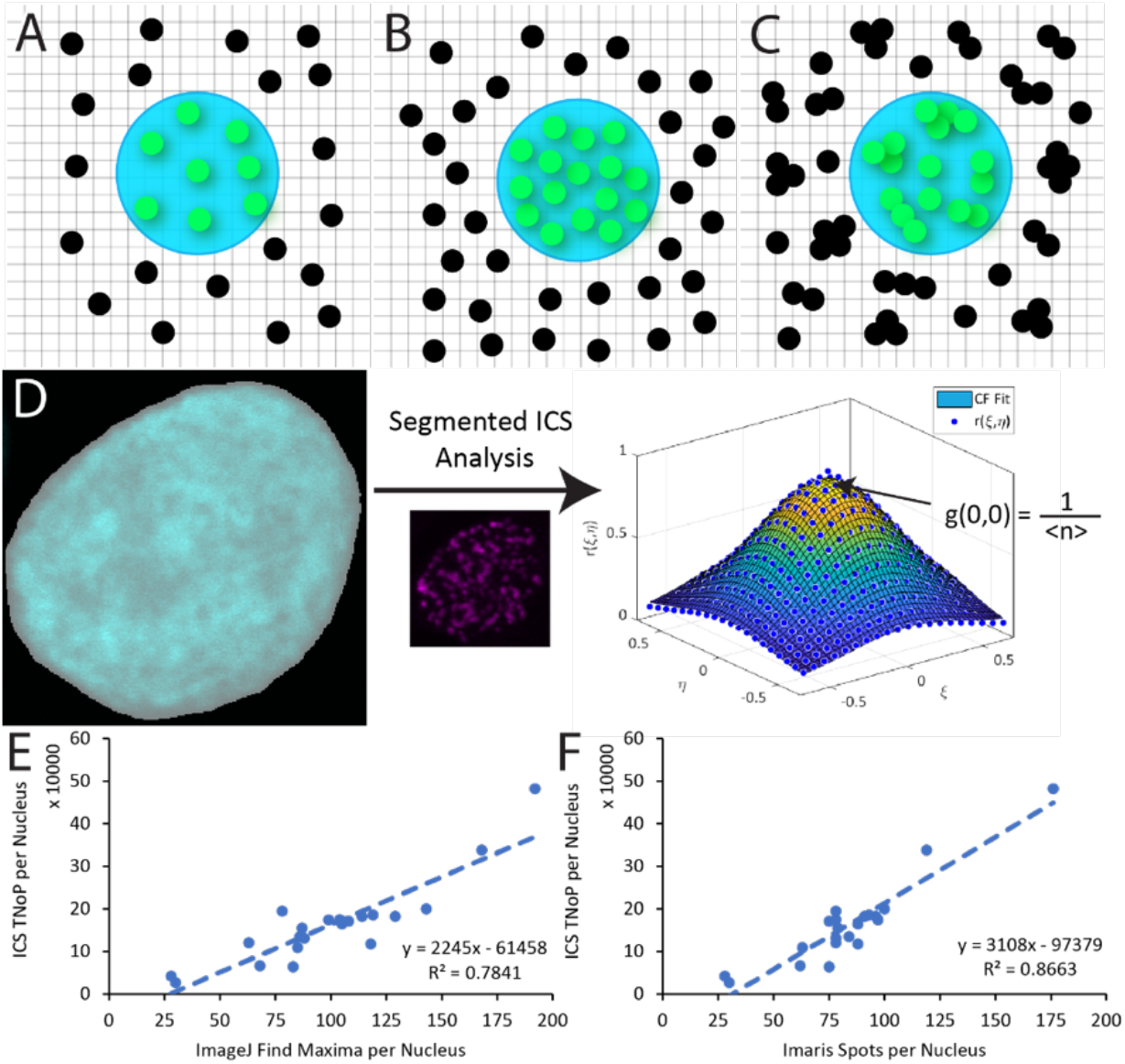
Counting foci using image correlation spectroscopy correlates with optimized measurements using existing approaches. (**A**) Schematic population of monomeric fluorescent particles where emitting particles (green) are in a diffraction limited excitation focus (cyan) – not to scale. (**B**) Schematic showing increase in particle density from A) which results in an increase in average fluorescence intensity. (**C**) Schematic showing clustering of fluorescent particles where the total number of independent particle clusters is the same as A). Spatial ICS measures the average number of independent fluorescent units per area which, along with the average fluorescence intensity, is used to calculate the average degree of aggregation of the particles. (**D**) Intensity-based segmentation of the DAPI channel with an automatic threshold using the MATLAB function *imbinarize* to create a mask of the nucelus. The mask specifies the ROI for ICS analysis of the γH2AX channel, outputting mean number of independent fluorescent particles per focal spot area (inverse of the intensity normalized spatial correlation function amplitude). (**E**) and (**F**) Comparison of the ICS parameter TNoP with E) ImageJ Find Maxima tool with a user defined threshold and F) Imaris Spots with an automatic threshold for the same nuclei. A linear line of best fit is shown on each graph with the resulting R^2^.

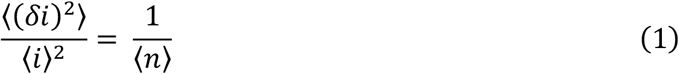

given intensity fluctuations defined as *δi* = *i* − ⟨*i*⟩, where ⟨*i*⟩ is the mean intensity and ⟨*n*⟩ is the mean number of independent fluorescent particles per focal spot area.

To obtain this number density information independent of white noise sources (such as shot noise), ICS calculates the mean intensity normalized spatial autocorrelation function of fluorescence intensity fluctuations (Eq. 2) of a region of interest (ROI). The orthogonal spatial lag variables *ξ* and η represent discrete pixel shifts in *x* and *y* directions in an image at which the spatial correlation is calculated. The zero spatial lags value of the spatial autocorrelation function is the square relative intensity and hence the particle number density.

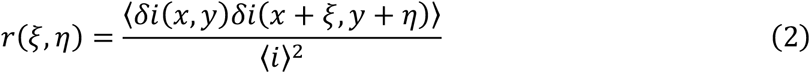

For computational speed, the spatial correlation function is calculated using Fourier methods (Eq. 3)^28^, where *F* is the discrete 2D spatial fast Fourier transform of the ROI, *F\** is the complex conjugate and *F*^*-1*^ is the inverse Fourier transform.

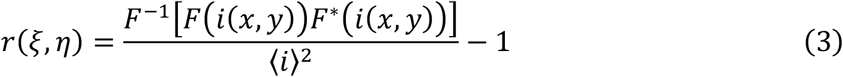

The calculated spatial intensity fluctuation correlation functions are then each fit to a 2D Gaussian (Eq. 4) using a non-linear least-squares algorithm, where the zero lags point is not weighted due to white noise contributions. Output fit parameters are ***g*(0, 0)**, the zero-lags amplitude, ***ω***_**0**_, the e^-2^ Gaussian correlation radius, and ***g***_∞_, the long spatial lag offset.

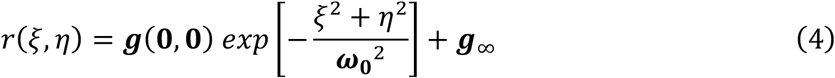

The best fit zero lags amplitude of the correlation function is an estimate of the square relative fluctuation from Eq. 1, with an inverse that is the mean number of independent fluorescent particles per focal spot area, ⟨*n*⟩ (Eq. 5). The total number of particles (TNoP) in the ROI can be calculated using the image area and ***ω***_**0**_ (Eq. 6).

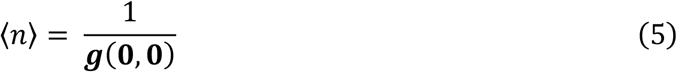

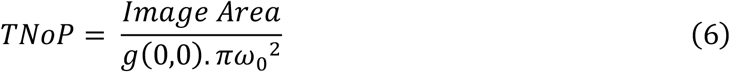

When fluorescent molecules undergo aggregation or clustering, the clusters/aggregates are not resolvable using diffraction limited optical microscopy (i.e., confocal) unless large aggregates are present (such as individually distinguishable foci). However, aggregation manifests in the ROI as larger relative intensity fluctuations from a smaller number of brighter fluorescent particles per focal spot. If the number of overall fluorophores does not change, the mean fluorescent intensity should be constant, but the number of independent fluorescent particles (formed of one or more fluorophores) should decrease after clustering (**Fig. 2 B-C**). Since the average intensity of the ROI is proportional to the total number of fluorophores ⟨*n*_***f***_⟩ after background correction and a constant (c) relating intensity to fluorophore count, a degree of aggregation (DA) measurement can be calculated using the average intensity, ⟨*i*⟩, and the mean number of fluorescent particles, ⟨*n*⟩;

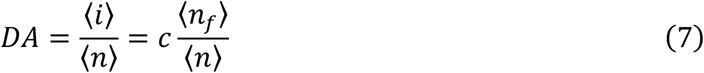

To first order, the degree of aggregation is proportional to the mean number of fluorophores per aggregate if the variance of the aggregate distribution is small^29^.

The output from ICS calculations can be used to understand the distribution of fluorophores contributing to the image. The two outputs used here are TNoP and DA. TNoP is a measure of the inverse of the number of independent particles – this calculation is not dependent on the size (or number of fluorophores) contributing to each independent particle, nor the distribution of particle sizes. The DA captures the size of each independent cluster through dividing the average voxel intensity by the average number of independent particles per voxel. Thus, the DA captures how many fluorophores comprise each independent particle in the image.

### Image Analysis

For the ICS analysis, nuclei were segmented from the DAPI channel using a watershed approach in MATLAB. Briefly, we applied a difference of gaussians for size-based feature extraction of nuclei in the image (upper gaussian σ=100, lower gaussian σ=2-5 dependent on cell size). This feature enhanced image was then binarized to create a primary mask (using the function *imbinarize*) and connected components under 100 pixels were removed (using the function *bwareaopen*). We then created a distance transform of the resulting binary image (using the function *bwdist*). Seed points were determined by finding the regional maximum of a gaussian filtered image, with a gaussian σ of appropriate size for seed point detection (σ=7 or 8 dependent on cell size). The seed points were applied to the distance transform, and we applied the *watershed* function to the resulting image to obtain masks for each nucleus (extracted using the function *regionprops*).

Using the nucleus masks, we performed ICS analysis on the antibody-stained channel, within size limitations for a nucleus (objects > 400 and < 2400 pixels). Intensity, DA and TNoP were calculated for each cell. In this analysis, since the ROI is defined by the nucleus mask (segmented from nuclear signal using DAPI staining), TNoP represents the total number of particles per nucleus, with the image area being the number of pixels in each nuclear mask.

Measurements across experiments were normalized by the control in each dataset. The confocal imaging settings were kept the same across each dataset to provide comparable measurements per dataset. Data were plotted in Prism and statistical analysis was performed in Excel.

Local maxima were used to count foci in ImageJ/FIJI, using the tool *Find Maxima* under the *Process* menu^31^. This tool requires a threshold input parameter (Prominence/Tolerance), where a threshold is set at the maximum value minus noise tolerance and local maxima must be greater than this threshold to be counted. To compare the effect of the input parameter, we performed measurements at Tolerance = 300 and 600. The final measurement was taken at the user-defined threshold where the user visually inspected different Tolerance values and chose a value that visually removed background (thresholds between 300 and 600) – a subjective process.

Imaris spots analysis were performed with Imaris 9.8 with background subtraction, using a spot size of 1 µm that was measured from the smallest distinguishable foci from the control image. For images with multiple nuclei, ImarisCell was used with the detection type “cell and vesicles”, where the nucleus in the DAPI channel was modelled as the “cell” and antibody-labeled foci were modelled as the “vesicles”. This enabled us to count the number of foci per nucleus for multiple cells. In each dataset, the automatic “quality” threshold was determined from the control image and the same threshold was used for every image in the dataset to be able to compare the outputs across datasets. Similar to the ICS analysis, only cells with an area between 400-2400 voxels were included for analysis.

### Materials and Cell Culture

All chemicals were purchased from Sigma unless otherwise noted. All drugs were purchased from Selleck Chemicals. Cells were cultured in media supplemented with 10% fetal bovine serum and 1% penicillin/streptomycin. OVCA429 were originally obtained from Dr. Michael Birrer, HT1080 were obtained from ATCC cultured in DMEM. SKOV3, HCC1937, HCC1395, MCF7, UWB1.289 and UWB1.289 +BRCA1 were obtained from ATCC and cultured in RPMI. UWB1.289res were made resistant to olaparib through long-term culture in up to 1 μM olaparib. Unless otherwise indicated, for MMS treatment, cells were incubated for 6 hours before fixation with 4% formaldehyde, whereas for treatment with PARP inhibitors and other drugs, cells were incubated for 24 hours before fixation.

### Cell Dose Dependence Response

Cells were plated in a 96-well plate at 2×10^3^ cells per well and treated with drug for 5 days in five replicate wells. Cell viability was then determined using Presto Blue (Thermo Fisher). Signal was averaged over replicate wells and normalized to blank (no cells) as well as to the levels of untreated wells. A sigmoidal response curve was fit to the average of two experiments in Prism.

### Immunofluorescence and Fluorescence Microscopy

Cells were plated in 8-well slides (Fisher) or 384 well glass bottom plates (CellVis) to achieve 50% confluence overnight. Cells were then treated and fixed with 4% formaldehyde in PBS at room temperature for 10 minutes. Antibody staining was performed according to the manufacturers protocol (Cell Signaling Technologies or Abcam). Primary antibodies, that were vendor validated for specificity, were γH2AX (CST, #2577), RPA1 (CST, #2267), RAD51 (Abcam, 63801) and PCNA (CST, #2586), with Anti-rabbit IgG Fab2 Alexa Fluor 647 (CST, #4414) as the secondary fluorescent antibody. After antibody labeling, prolong gold with DAPI (CST) or DAPI (1 μg/ml) in PBS for well plate experiments was added and samples were stored for up to 1 week prior to imaging. Confocal imaging was performed on an Olympus FV1000 multiphoton/confocal microscope using an Olympus XLUMPlanFL N 20x objective, NA 1.00 with chromatic correction, pinhole at 1 Airy unit and 3x digital zoom. DAPI was excited with a 405 nm laser, while Alexa Fluor 647 was excited with a 635 nm laser. A multiband (DM 405/488/561/633 nm) dichroic mirror was used to direct the excitation laser and collect emission. Emission was then separated into two channels through a dichroic mirror (490nm) and DAPI signal was cleaned up by diffraction grating, which is the low band pass filter used by this instrument. Widefield imaging were performed on a Nikon Ti2 with a 20x (NA 0.75) objective using standard DAPI and Cy5 filter cubes.

### Western Blotting

Cells were lysed with RIPA buffer (Thermo Fisher) containing protease inhibitors (Thermo Fisher), vigorously vortexed and incubated on ice for 15 minutes. Lysate was then pelleted by centrifugation at 14,000g for 15 min at 4 ºC. Supernatant was transferred to a clean tube and total protein was determined using a BCA assay (Pierce). Protein was loaded into a NuPAGE gel (Thermo Fisher) and transferred to nitrocellulose paper (Thermo Fisher). Blocking and antibody labeling was then carried out according to the antibody manufacturer’s protocol. GAPDH (R&D systems, #AF5718) was used as a loading control.

### Statistics and Reproducibility

All experiments were performed in multiple repeats as indicated in the figure legends. Normalization to control was performed for conditions during each imaging session by dividing each datapoint by the average of the control data and the average control value was set to 1. For results that were collected from multiple different experiments the data were normalize to the control for each experiment. Statistical significance was calculated compared to the next lower dose using one way ANOVA or Students’ t test.

## Results

### Counting Foci via Fluctuation Analysis

Most foci counting methods are based on finding the local fluorescence intensity maxima either through intensity thresholding to create masks of high intensity single foci or implementing local maxima finding algorithms. Here, we evaluated traditional local maxima finding algorithms on the ovarian cancer cell line SKOV3 by creating resolvable DNA damage with the PARP inhibitor olaparib (10 µM) and antibody-labeling γH2AX to visualize foci (**Fig. 1B**). To enable clean isolation of nuclei, the cells were also labeled with DAPI (**Fig. 1A**). First, we used the Find Maxima tool in ImageJ/FIJI which detects local maxima to define individual foci. However, this approach relies heavily on the intensity threshold parameter (Prominence/Tolerance) used to remove the background signal. This parameter is defined by the user and is highly subjective – a high value causes some foci to be omitted from the count, yet, conversely, a low value leads to background signal being counted as foci. Therefore, we evaluated images at 2 different intensity thresholds (**Fig. 1D-E**) in 21 single cells to determine the ambiguity of the threshold parameter (**Fig. 1G**) before manually determining a threshold for each cell for the final measurement (**Fig. 1H**).

In addition, we implemented a different local maximum finding algorithm with automatic thresholding using Imaris Spots detection (**Fig. 1F**). Spots detection uses a Gaussian filter to smooth the image and foci are then located at the local maxima of the filtered image. Automatic thresholding is based on a k-means statistical method and Spots uses an expected foci size to identify multiple spots in a large cluster where individual foci are indistinguishable and could be classified as single foci with local maxima detection. Our results produce foci counts that are dependent on the approach used and setting implemented (**Fig. 1 and S1**). In our analysis, different thresholds/algorithms give different counts for the same cell due to different intensity foci (white arrows, **Fig. 1I**), different size foci (green arrows, **Fig. 1I**) and non-uniform background intensities (blue arrows, **Fig. 1I**).

We next employed fluctuation analysis via spatial ICS in the same set of images. For all ICS measurements immuno-staining was performed with primary and secondary antibodies based on the vendor protocol. No pre-extraction was performed prior to fixation. ICS was applied to individual, segmented nuclei in automated scripts. Uniquely ICS analyzes the entire dynamic range of detected fluorescence, not just intensity peaks that could be found with local maxima algorithms. We employed ICS analysis on segmented nuclei (**Fig. 2D**) and calculated the total number of particles (TNoP) parameter per nucleus for comparison to foci counting data from ImageJ/FIJI and Imaris. Here, TNoP is not a measure of foci, rather the amount of non-diffuse signal. Direct, cell by cell comparison of ICS TNoP with both the user-optimized threshold ImageJ Find Maxima (**Fig. 2E**) and Imaris Spots (**Fig. 2F**) foci counting results produced a strong correlation. Here, ICS is determining the total number of independent particles, leading to a TNoP value that is not directly counting resolvable foci. However, the correlation to the other approaches demonstates that ICS is measuring traditional foci forming DNA damage.

### Particle Counting versus Aggregation in γH2AX signal

One major goal of foci counting in cells is to determine the degree of DNA damage. Therefore, after validating our foci counting methods at the individual cell level, we sought to evaluate ICS as a tool to quantify the DDR response in bulk populations of cells. We first treated SKOV3 cells with varying concentrations of methyl methane-sulfonate (MMS) or the PARP inhibitors veliparib or olaparib and evaluated DNA damage through immunofluorescence of γH2AX. MMS is a DNA alkylating agent that directly damages DNA by modifying guanine and adenine to instigate mispairing of bases and replication blocks^32^. PARP inhibitors, however, do not necessarily directly induce DNA damage, but stall the DNA damage response by “trapping” PARP1 and preventing recruitment of critical DNA damage response proteins^23,33-35^. Therefore, these two drug classes should have different impacts on focus formation and focus composition.

MMS treatment produces a wide range of γH2AX intensity and visible foci across individual cells (**Fig. 3A-C**; 0.1 mM MMS and **Fig. S2**; 0 mM and 1 mM MMS). Thus, analysis capable of detecting both low intensity foci and overlapping, highly abundant foci within a single experiment is needed for accurate measurements over a range of conditions. In the non-contrast adjusted γH2AX image (**Fig. 3B**), brighter γH2AX foci are clearly distinguishable, however once the image is contrast adjusted (**Fig. 3C**), lower intensity foci are apparent and the higher intensity foci appear as large foci clusters, especially in nuclei demonstrating more DNA damage. To overcome the inherent distribution of DNA damage across cells, we scaled up the analysis to examine hundreds of cells per image by acquiring large, stitched images (**Fig. S3**). Here, due to the dependency of selecting the proper background intensity per cell, we omitted the ImageJ Find Maxima analysis and analyzed data using ICS and Imaris Spots. To directly compare analyses, each measurement within an experiment was normalized to the average value of DMSO treated control cells.

**Figure 3.**
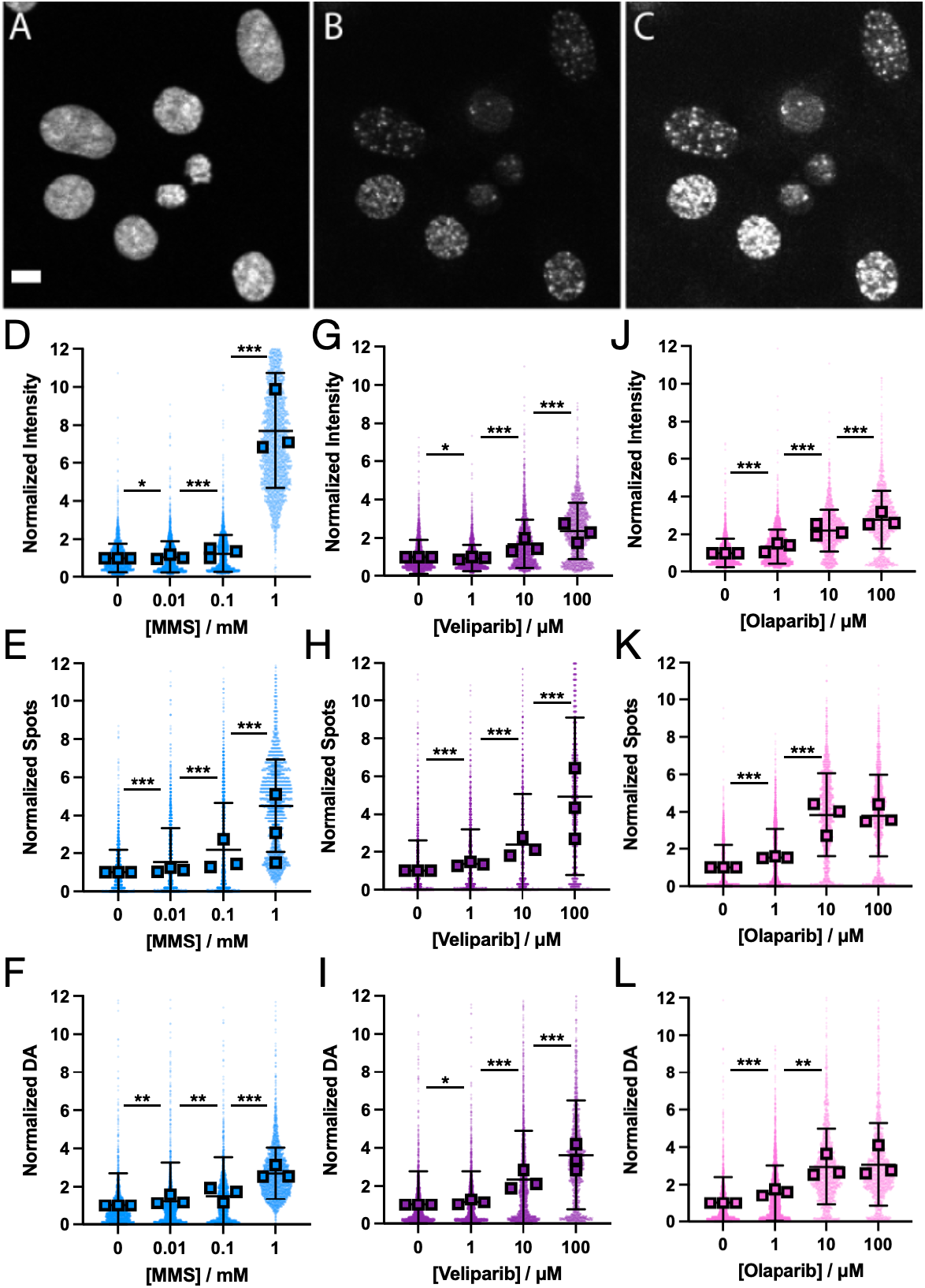
Imaris Spots and ICS capture dose dependent DNA damage through γH2AX foci in SKOV3 cells. (**A-C**) Confocal images of SKOV3 cells treated with 0.1 mM MMS; A) DAPI channel with 10 µm scale bar, B) γH2AX channel, C) γH2AX channel with intensity-adjustment. (**D-L**) Data for normalized intensity, normalized number of spots and normalized ICS DA of γH2AX signal across different MMS (**D-F**), veliparib (**G-I**) and olaparib (**J-L**) treatment concentrations in SKOV3 cells. Shown are individual cells, average for each biological repeat (n=3, squares), and average with standard deviation. Results are normalized to average DMSO. MMS; N=2423-3160, veliparib; N=1818-2922, olaparib; N=1489-2956. * indicates p<0.05, ** indicates p<0.001, *** indicates p<1×10^−10^ via one-way ANOVA significance testing.

In analyzing DNA damage, the amount of γH2AX per cell is represented by the normalized integrated fluorescence intensity (**Fig. 3**). We found that γH2AX intensity experienced small, significant increases at lower MMS doses but showed a massive (nearly 8X), significant increase at the highest dose, results that agree with SKOV3 sensitivity to MMS (**Fig. S4**). PARP inhibitor treatment produced similar results except for a slight decrease at 1 µM veliparib. The number of spots determined by Imaris captures the dose dependent DNA damage induced by MMS and PARP inhibitors as expected (**Fig. 3**), with the response to olaparib plateauing after 10 µM. Yet, surprisingly, at the highest dose of PARP inhibitor, the number of foci was higher under veliparib treatment, the weaker PARP inhibitor (**Fig. S4**).

We then analyzed the same images with ICS. Microscopic number density calculated by ICS often decreases in cases of significant aggregation, resulting in a smaller number of more densely packed clusters^36,37^, therefore we calculated the ICS degree of aggregation (DA) as a measurement that captures both low and high DNA damage. Critically, ICS DA is not a direct measurement of the number of foci, but rather a quantification of overall label clustering. DA results demonstrated both MMS and PARP inhibitor dose dependence (**Fig. 3**), similar to results obtained with Imaris Spots. Like Spots analysis, the ICS DA analysis of 100 µM veliparib produced a higher value than 100 µM olaparib, compared to their respective controls. Here, ICS DA proved comparable to Imaris Spots.

We next analyzed DNA damage dose dependence in another cell line, OVCA429. Fig. 4A-C displays OVCA429 cells exposed to 0.1 mM MMS, highlighting a range of γH2AX signal, spanning from only low diffuse signal in the nucleus, to single distinguishable γH2AX foci in the diffuse signal, to large clusters of different sizes in high intensity nuclei (**Fig. 4C**). γH2AX intensity significantly increases at 1 mM MMS (**Fig. 4D**) and Spots analysis captured the MMS dose dependent DNA damage (**Fig. 4E**) and ICS DA produced results similar to Spots analysis (**Fig. 4F**).

**Figure 4.**
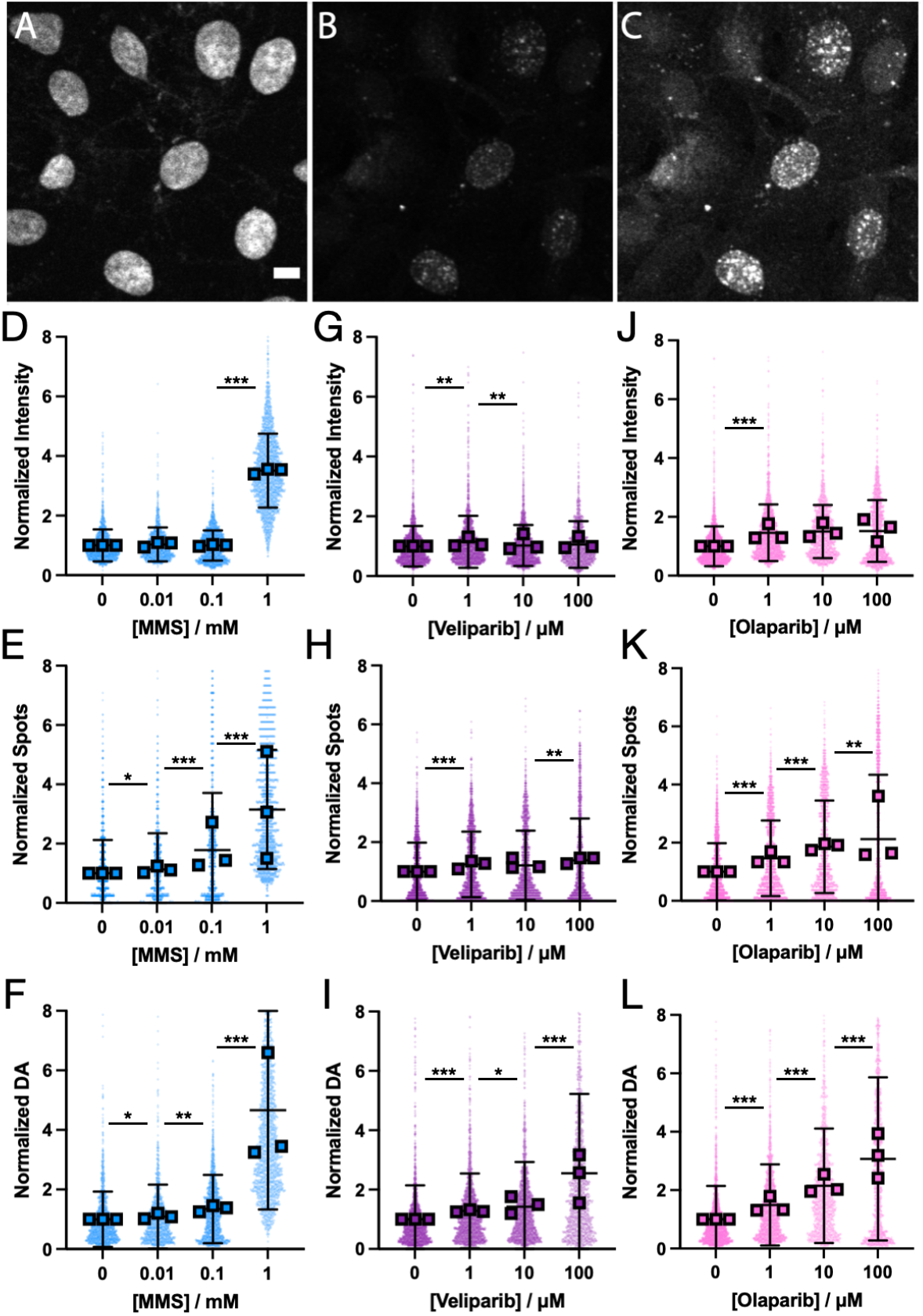
Imaris Spots and ICS DA capture dose dependent DNA damage through γH2AX in OVCA429 cells. **A-C**) Confocal images of OVCA429 cells treated with 0.1 mM MMS; A) DAPI channel with 10 µm scale bar, B) γH2AX channel, C) γH2AX channel with intensity-adjustment. (**D-L**) Data for normalized intensity, normalized number of spots and normalized ICS DA of γH2AX signal across different MMS (**D-F**), veliparib (**G-I**) and olaparib (**J-L**) treatment concentrations in OVCA429 cells. Shown are individual cells, average for each biological repeat (n=3, squares), and average with standard deviation. Results are normalized to average DMSO. MMS; N=1194-2068, veliparib; N=1256-1997, olaparib; N=1445-2334. * indicates p<0.05, ** indicates p<0.001, *** indicates p<1×10^−10^ via one-way ANOVA significance testing.

Contrary to SKOV3 analysis, despite similar sensitivity to olaparib and veliparib (**Fig. S4**), γH2AX intensity in OVC429 in response to PARP inhibitors showed a lower change in signal with no significant change in intensity at higher concentrations (**Fig. 4**). However, Imaris Spots captured a significant dose-dependent response with all but 10 μM veliparib generating a significant increase from the next lower dose (**Fig. 4**). Thus, despite no changes in intensity, more foci are being formed at higher doses. ICS analysis, however, was more sensitive than Imaris Spots, capturing a significant dose-dependent change at each concentration (**Fig. 4**). Overall, in OVCA429, ICS DA produced similar results to Imaris Spots in olaparib treated cells, but measured dose dependent response in cells treated with veliparib where Imaris Spots was unable to capture the dose dependency.

### ICS Captures Clustering of DNA Damage Proteins that do not form Resolvable Foci

Next, we sought to quantify the behavior of proteins that do not form resolvable foci during DNA damage without immunofluorescence pre-extraction, specifically Replication Protein A1 (RPA1) and RAD51. The RPA complex is required for major DNA repair pathways and modulates RAD51 recruitment to sites of DNA damage^38,39^. However, increased RPA1 and RAD51 concentrations created at sites of DNA repair are not always resolvable as individual foci^39,40^. In the absence of induced DNA damage, RPA1 immunofluorescence produces a diffuse signal throughout the nuclei in SKOV3 cells (**Fig. 5**). Only with contrast adjusting (**Fig. 5C**), does heterogeneous distribution of RPA1 become visible.

**Figure 5.**
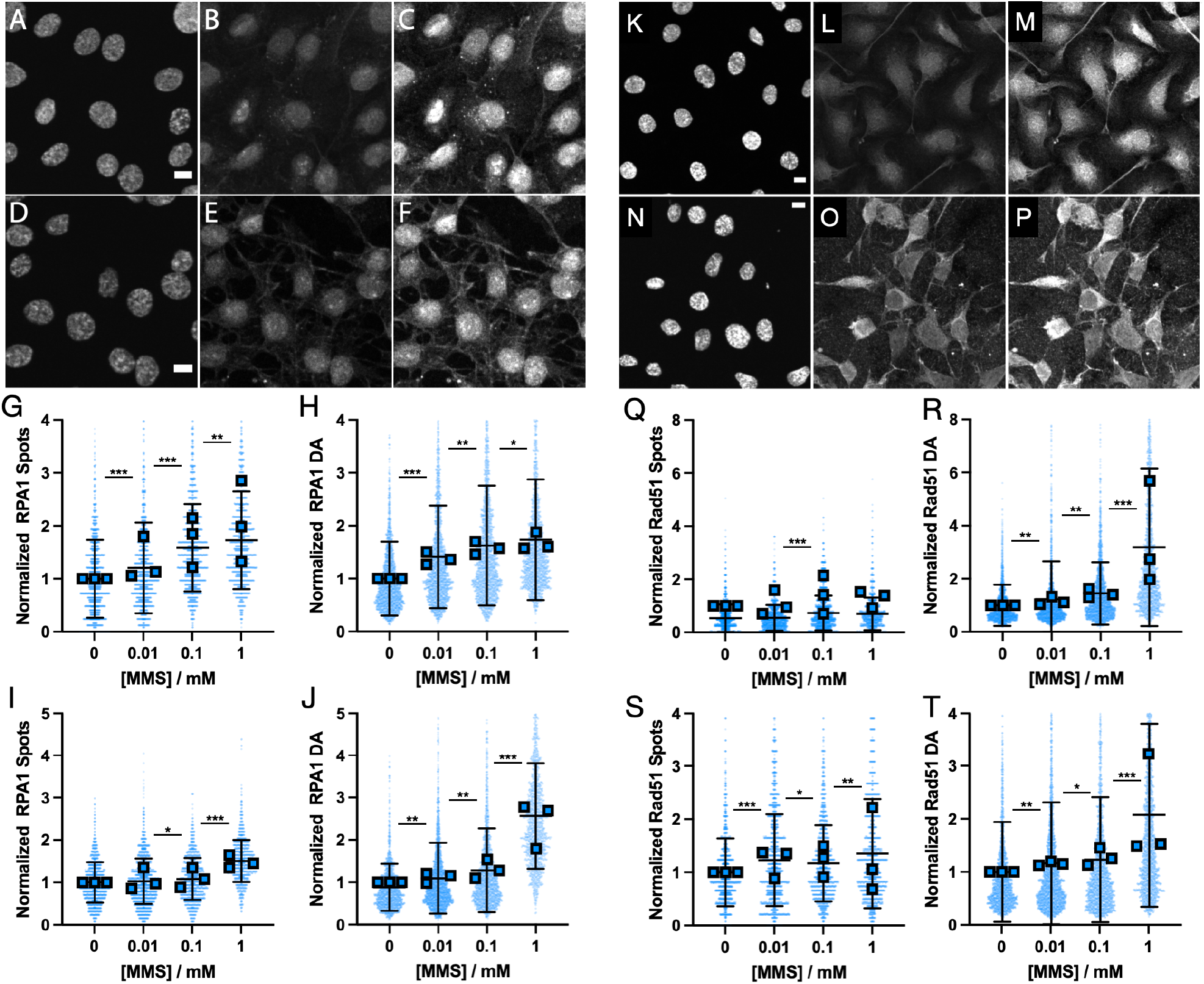
Dose-dependent DNA damage impact on RPA1 and RAD51. (**A-C**) Confocal images of control SKOV3 cells, (**D-F**) SKOV3 cells treated with 1 mM MMS; (**A&D**) DAPI channel with 10 µm scale bar, (**B&E**) RPA1 channel, (**C&F**) RPA1 channel with intensity-adjustment. Normalized RPA1 Imaris Spots (**G**) and normalized ICS DA (**H**) in SKOV3 and OVCA429 cells (**I&J**). Shown are individual cells, average for each biological repeat (n=3, squares), and average with standard deviation. Results are normalized to average DMSO. N=1600-1872 for SKOV3 and N=1320-2198 for OVCA429. (**K-M**) Confocal images of control SKOV3 cells, (**N-P**) SKOV3 cells treated with 1 mM MMS; (**K&N**) DAPI channel with 10 µm scale bar, (**L&O**) RAD51 channel, (**M&P**) RAD51 channel with intensity-adjustment. Normalized RAD51 Imaris Spots (**Q**) and normalized ICS DA (**R**) in SKOV3 and OVCA429 cells (**S&T**). Shown are individual cells, average for each biological repeat (n=3, squares), and average with standard deviation. Results are normalized to average DMSO. N=1597-2241 for SKOV3 and N=1822-2311 for OVCA429. All results, * indicates p<0.05, ** indicates p<0.001, *** indicates p<1×10^−10^ via one-way ANOVA significance testing.

We used MMS to induce DNA damage in SKOV3 and OVCA429 cells. As expected, normalized intensity for RPA1 and RAD51 values (**Fig. S5**) did not display a dose dependence on MMS concentration. In SKOV3 cells treated with 1 mM MMS (**Fig. 5**), RPA1 recruitment to sites of DNA damage is observed alongside the diffuse signal of the nuclei. We first quantified the dose-dependent distribution of RPA1 with both Imaris Spots analysis (**Fig. 5**) and ICS DA measurements (**Fig. 5**). In SKOV3 cells, both Spots analysis (**Fig. 5G**) and ICS DA analysis (**Fig. 5H**) captured a dose dependent increase in RPA1 clustering, with the repeatability between experiments being much higher in ICS DA analysis suggesting that differences in immunofluorescence labeling between experiments impact Spots more than DA. In OVCA429 cells, Spots analysis demonstrated a more modest increase with increasing MMS dose (**Fig. 5I**), however, DA analysis produced a significant, larger increase in aggregation exceeding 2.5x of control at 1 mM MMS (**Fig. 5J**). Analysis of γH2AX clustering in OVCA429 cells treated with MMS (**Fig. 4G**) demonstrates a large increase at 1 mM MMS, suggesting that RPA1 DA analysis (**Fig. 5J**) is capturing DNA damage with a similar sensitivity to γH2AX analysis. The increased sensitivity in ICS DA versus Spots also indicates that measuring aggregation instead of foci counting could be more sensitive when analyzing proteins that do not form distinct foci.

Similar to RPA1, RAD51 does not form distinguishable foci in the nuclei in cells without pre-extraction (**Fig. 5**). With MMS treatment, Spots analysis of RAD51 foci captured an MMS dose dependency in OVCA429, but not SKOV3 cells (**Fig. 5**). However, ICS DA analysis captured dose dependent increases in aggregation (**Fig. 5**), with a 3X increase over control in SKOV3 at 1 mM MMS. Similar to RPA1 labeling experiments, the MMS dose dependence observed with RAD51 DA analysis is comparable to trends observed with γH2AX DA analysis for both cell lines. Thus, ICS DA proved to be more sensitive to RPA1 and RAD51 aggregation in response to DNA damage that induces significant γH2AX foci formation.

We then sought to determine if ICS analysis could provide insight into the DDR that had been previously unknown. Surprisingly, when we measured RPA1 DA induced by PARPi treatment we found different trends in different cell lines (**Fig. 6**). HCC1937 cells show a positive dependency of RPA1 DA with olaparib concentration. However, HT1080 cells show the opposite relationship, with RPA1 DA decreasing with increasing olaparib concentration. We then expanded our analysis to more cell lines and measured RPA1 DA, normalized to control, in response to 1 μM treatment of three different PARPi (**Fig. 6B**). Plotting the RPA1 DA value for each PARPi as a function of the cellular sensitivity (cell IC_50_) to each drug generated a curious pattern (**Fig. 6C**). Here, cell lines that were most sensitive to PARPi treatment demonstrated a decreasing RPA1 DA with drug potency (inverse of cell IC_50_), while cell lines that were more resistance had the opposite response. Fitting a linear curve to the RPA1 DA vs. cell IC50 data for each cell line and plotting the slope of that fit against the average cell line sensitivity to PARPi treatment produced a striking pattern (**Fig. 6D**). These results suggest the capacity to recruit RPA1 into the DDR imparts resistance to PARPi treatment. Surprisingly, the expression level of RPA1 in each cell line had no correlation to cell line sensitivity to PARPi (**Fig. 6E&F**). Therefore, these results demonstrate the power of ICS to identify DDR responses that would be otherwise very difficult to detect.

**Figure 6.**
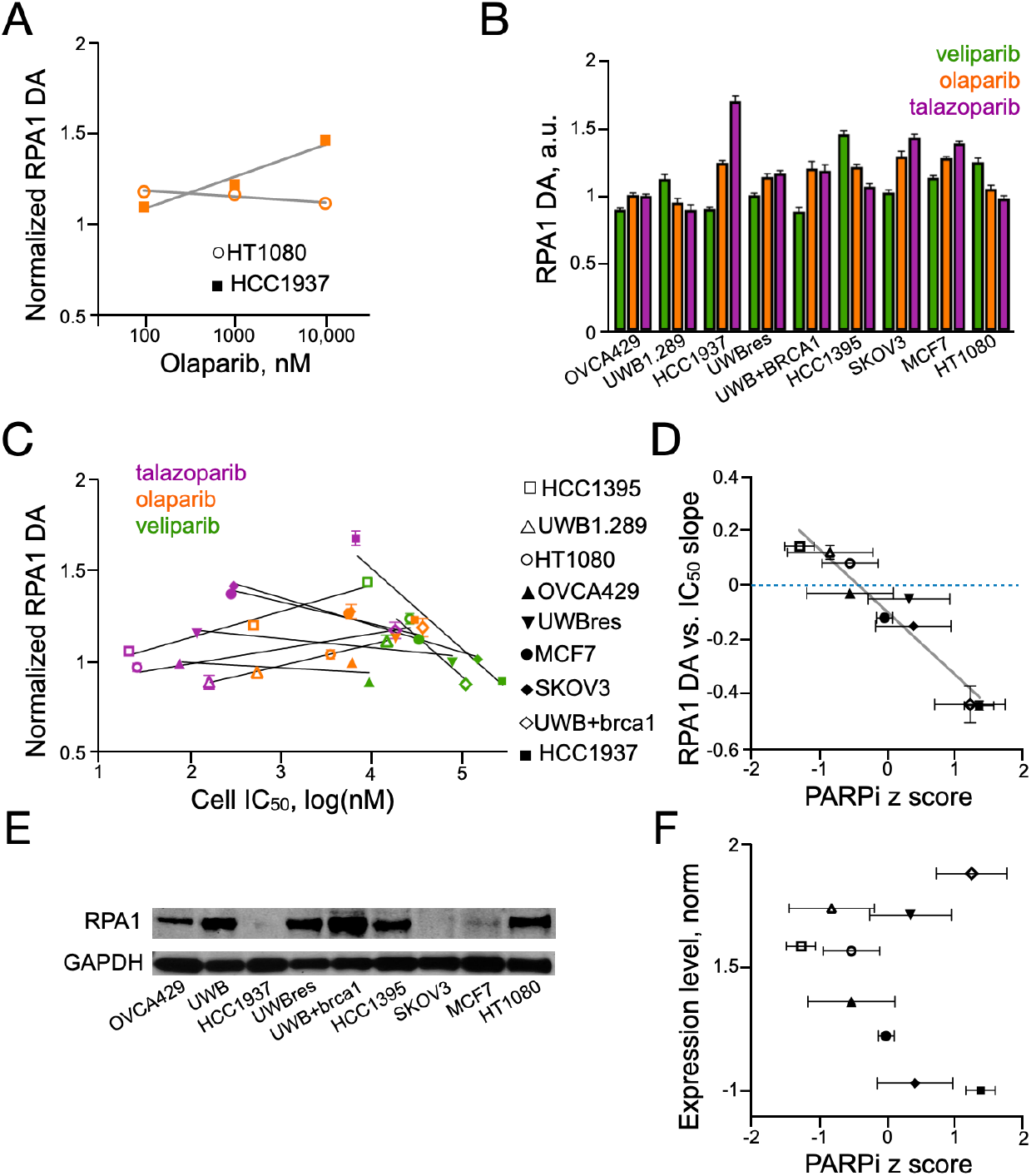
RPA1 recruitment correlates with cell sensitivity to PARPi. (**A**) RPA1 DA as a function of olaparib concentration for HT1080 and HCC1937 cells, shown are average with SEM, n >= 504 cells, 3 biological repeats, and linear fit. (**B**) RPA1 DA in cells treated with 1 μM PARPi overnight, shown are average, normalized to untreated control, with SEM, n >= 350 cells, 2 biological repeats. (**C**) RPA1 DA as a function of cell line IC50 for each cell line and PARPi shown in Fig. 5b. Shown are average, normalized to untreated control, with SEM, n >= 350 cells, 2 biological repeats and linear fit (prism) for each cell line. (**D**) The linear fit slope of RPA DA vs. cell line IC50 as a function of average PARPi z-score for each cell line. Shown are slope fit with SEM and average z-score over three PARPi with st. dev. (**E**) Western blot of RPA1 expression levels for each cell line, with GAPDH loading control. (**F**) Normalized protein expression level in each cell line as a function of PARPi z score. Shown are average z-score over three PARPi with st. dev.

### ICS can be Applied to a Diverse Set of Imaging and DNA Damage Response Measurements

Since ICS analysis is used on standard fluorescence microscopy images and we obtain a DAPI image every experiment we sought to incorporate cell cycle classification into measurements. Here, to demonstrate the broad applicability of ICS we performed measurements in 384 well plates on a widefield microscope. Previous results have established that DAPI intensity can be used to stage cells^41^. We confirmed our ability to distinguish cell stage through DAPI intensity by treating SKOV3 cells with palbociclib, camptothecin or etoposide, which cause cells to stall at different cell cycle stages. Similar to previous findings^41^, we found that palbociclib stalled cells in G1, camptothecin stalled cells in S and etoposide stalled cells in G2 (**Fig. 7A**). Once confirmed, we set DAPI intensity thresholds for each cell cycle stage. Thresholds were established automatically for each experiment by finding the peak of the DAPI intensity histogram of all control cells. We then classified G1 phase as cells with DAPI intensities within 10% of the DAPI intensity at the histogram peak, S phase as cells with DAPI intensity within 15% of the histogram peak times 1.5, and G2 phase as cells with DAPI intensity within 20% of the histogram peak times 2 (**Fig. 7B**).

**Figure 7.**
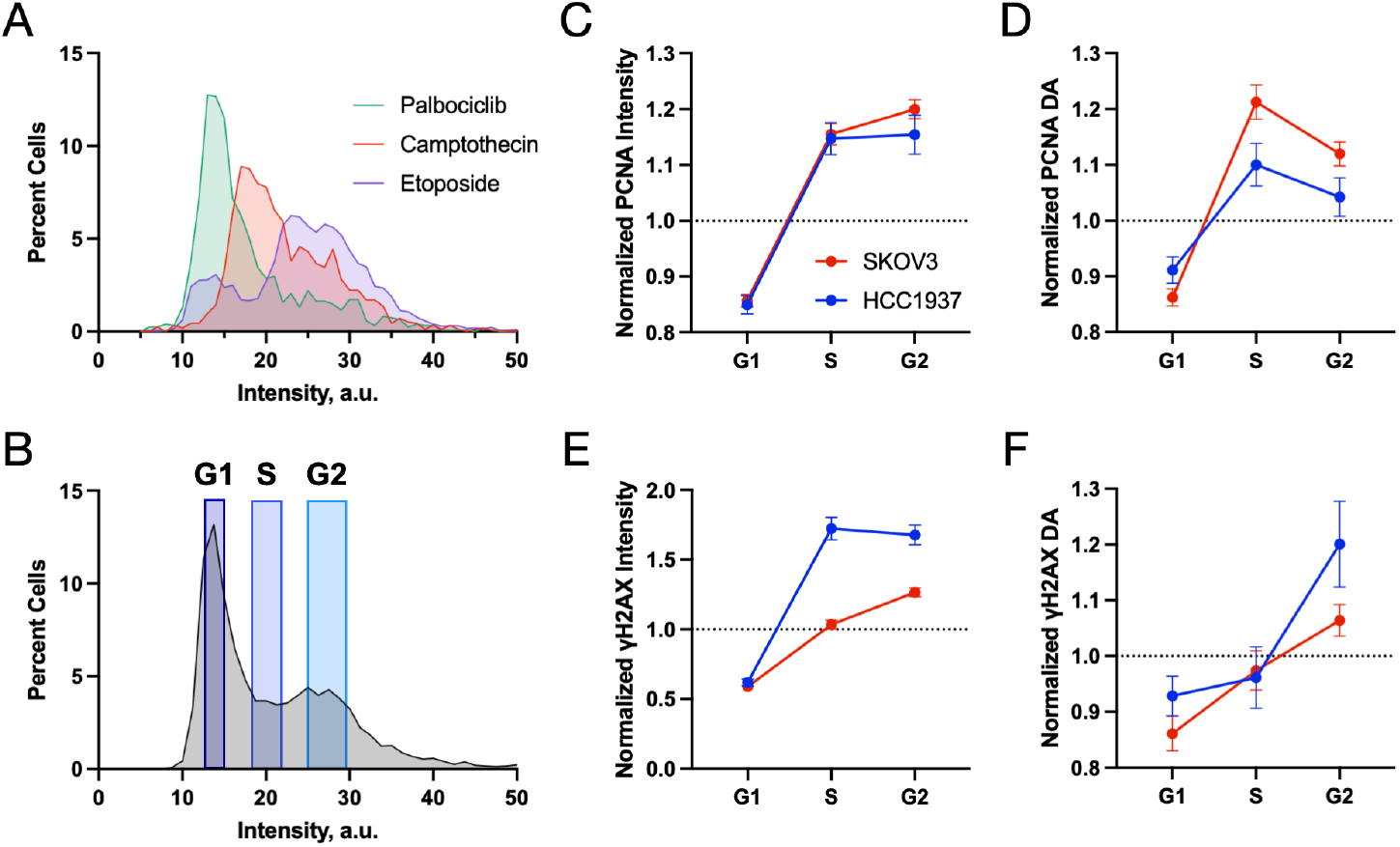
Incorporating and validating cell cycle data in widefield imaging. (**A**) Histograms of DAPI intensities for cells treated with palbociclib, camptothecin or etoposide, 1 μM overnight. (**B**) Representative assignment of cell cycle based on DAPI histogram (grey), using the first peak as G1 cells. Normalized PCNA intensity (**C**) and DA (**D**) in SKOV3 cells that had been classified by DAPI intensity into phases of the cell cycle. Normalized γH2AX intensity (**E**) and DA (**F**) in SKOV3 cells that had been classified by DAPI intensity into phases of the cell cycle. Shown are average n>200 cells per classification and standard deviation.

We validated our cell cycle staging approach through immunofluorescence of γH2AX and PCNA. Untreated SKOV3 and HCC1937 cells were classified by cell cycle stage and both ICS DA and intensity were calculated for the two labels. Data were normalized to unclassified average. We found that PCNA intensity increased from G1 to S, which was similar to G2 for both cell lines (**Fig. 7C**), suggesting that PCNA expression levels increase during S phase and plateau. However, PCNA DA increased from G1 to S, then decreased from S to G2 for both cell lines (**Fig. 7D**), suggesting that PCNA has the highest aggregation in S phase, as would be expected. PCNA is recruited to DNA during synthesis, which will increase the overall aggregation as measured here. We found that γH2AX intensity increased from G1 to S, then plateaued in G2 for HCC1937 cells but increased from S to G2 in SKOV3 cells (**Fig. 7E**). γH2AX DA increased from G1 to S and from S to G2 for both cell lines (**Fig. 7F**). These γH2AX results confirm that γH2AX is more prominent in S and G2 than G1.

We then measured γH2AX DA response in SKOV3 cells treated with DNA damaging agents olaparib, bleomycin or etoposide (**Fig. 8**). Cells were classified by cell cycle and the data aggregated and plotted as a function of drug concentration. Here we observed significant changes in both γH2AX DA and intensity as a function of damaging agent concentration. However, the γH2AX DA plateaued at higher doses, while the γH2AX intensity had a consistent dose dependent increase. These measurements suggest that the amount of γH2AX in the cell is a function of the degree of DNA damage, but the clustering of γH2AX reaches a peak at very large amounts of damage. Interestingly, we did not observe large differences between cell cycle stages for any of the compounds. Overall, olaparib and etoposide generated more γH2AX intensity in S and G2 phases while bleomycin generated the highest γH2AX intensity at 10 μM in G1 cells. However, the number of cells in each cell cycle was very dependent on the drug and concentration, with all compounds inducing more cells to stall at G2.

**Figure 8.**
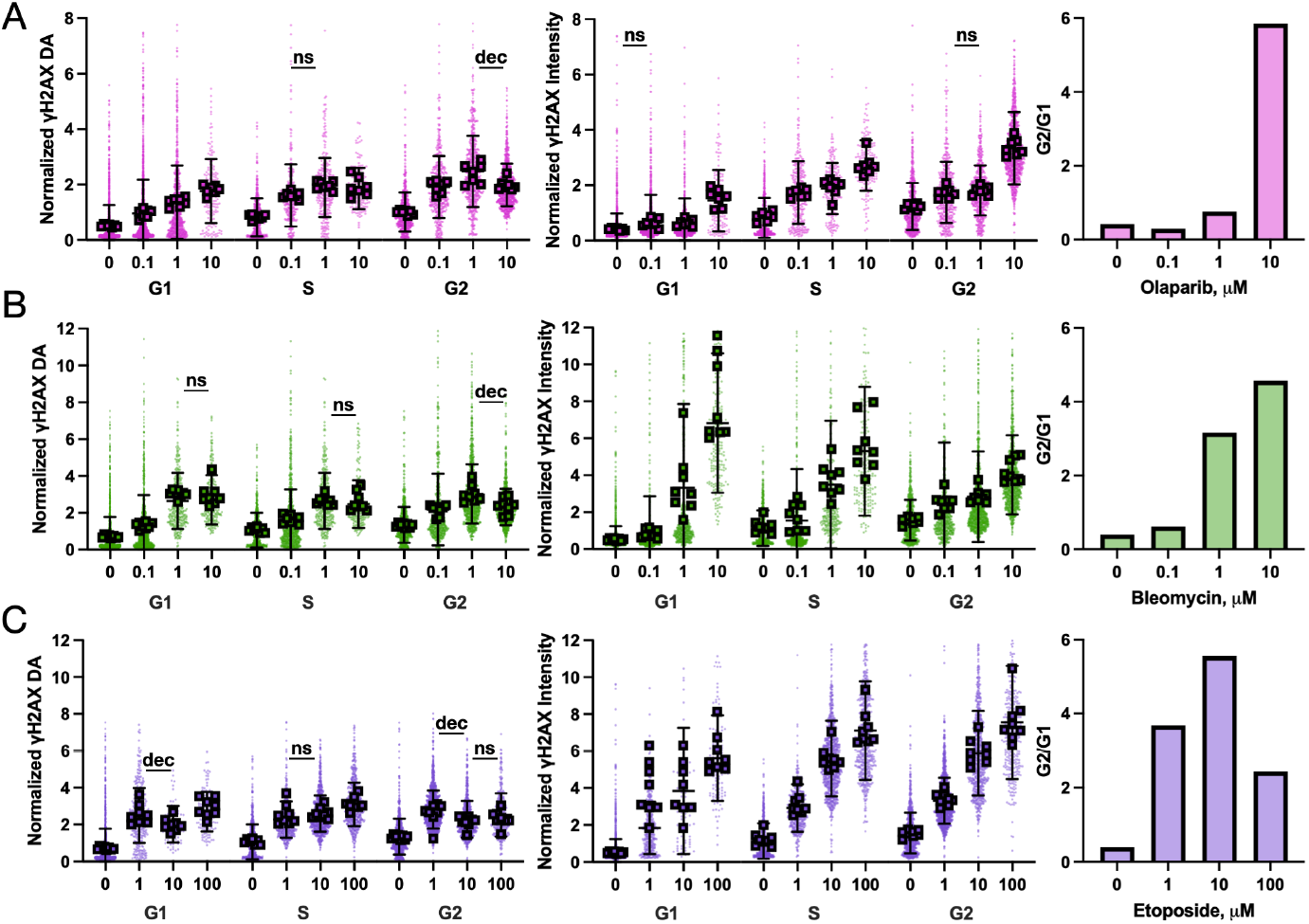
Cell cycle analysis of DDR ICS in widefield. Normalized γH2AX DA and intensity of SKOV3 cells treated with DMSO or olaparib (**A**), bleomycin (**B**) or etoposide (**C**) at 3 different doses (in μM) for 24 hours and separated by cell cycle stage based on DAPI intensity. Data were normalized to cell cycle independent DMSO average. Also shown are the ratios of the number of cells in G2 to G1 for each treatment condition. Data are n=7 biological repeats (box = average) with single cell values and standard deviation, n>200 cells per condition. For clarity all data are significantly different, p<0.05 Student’s t test from the next lower concentration unless marked – ns = no significance, dec = significant decrease.

## Discussion

Fundamentally, DDR foci are resolvable when the labeled protein concentration difference between foci and the rest of the nucleus is high enough to distinguish in fluorescence microscopy. The local concentration at sites of DNA damage is driven by either protein recruitment, or, in the case of γH2AX, post translational modifications^42^. Yet, most DDR proteins do not cluster in DNA damage at concentrations high enough to resolve recruitment in the absence of massive, artificial DNA damage. γH2AX is the most prominent marker of DNA damage foci largely because H2AX phosphorylation occurs at sites of double stranded breaks producing a marker with very low background signal. However, recent super resolution results identify γH2AX foci as groups of smaller foci^22^, suggesting that not all foci are equal, and large foci in diffraction-limited microscopy represent a greater degree of DNA damage than smaller foci. Therefore, foci count alone may not accurately represent DNA damage.

Clustering analysis with ICS overcomes both limits of counting traditional markers and measuring non-traditional markers as metrics of DNA damage or to study their role in the DDR. Initially we used the TNoP parameter to match foci counts in individual cells from two methods, however ICS calculated degree of aggregation (DA) captures overall clustering of protein and is therefore the more appropriate ICS measurement. Using ICS DA, we were able to detect the dose dependent response to two PARP inhibitors, matching or exceeding dose-dependent sensitivity of foci counting via Imaris Spots. Furthermore, ICS DA was uniquely able to measure dose-dependent DNA damage using RPA1 and RAD51 as DNA damage markers. The DNA damage response of RPA1 and RAD51 measured through immuno-staining have previously employed pre-extraction^38,43-45^, here the use of ICS removes this subjective sample preparation step to simplify measurements of the DDR activity of these proteins. The accuracy of these measurements was validated by the similarity of the dose dependency to γH2AX measurements. γH2AX is the most common marker used to define DDR foci. Therefore, ICS is able to evaluate DNA damage with non-traditional markers or evaluate the recruitment of proteins to DNA damage.

ICS analysis of γH2AX in OVCA429 cells treated with PARP inhibitors, where there was not a dose dependent increase of γH2AX intensity, revealed an increase in DA, indicating larger foci cluster formation was a merging of smaller clusters to produce an overall decrease in the number of foci clusters^36,37^. The formation of large foci can impact foci counting methods, however, Imaris Spots uses a spot size parameter that aids in placing multiple smaller spots per large foci. Yet, this spot size parameter failed to consistently capture the dose dependent increases in DNA damage where γH2AX intensity was not also increased. Spatial ICS is well suited to measure large aggregations and has previously been used to count distinguishable objects larger than the diffraction limit such as fluorescent beads^28^ and dendritic spines^46^, by using an intensity threshold to remove the background signal. This can be supplemented by various image analysis algorithms such as Gaussian filters to smooth out fluctuations in the objects so they can be detected as one object. However, we did not need an intensity threshold to remove the background signal, and only used a background subtraction that represented the intensity in a part of the image with no cells to account for autofluorescence outside of the cells.

We then used ICS to assess the involvement of RPA1 in the DDR, with results suggesting that recruitment correlates with cellular sensitivity to PARPi. These results were not apparent when measure RPA1 expression levels across cell lines, a typical evaluation of cell line sensitivity, indicating that ICS is capable of revealing underlying biology not detected by traditional approaches. We also demonstrated that cell cycle analysis can be readily performed on results to classify cells by cell stage for measurements of the impact of cellular stage on the DNA damage response. As a simple image analysis technique, ICS can be applied broadly to incorporate other analysis as well.

Here, we demonstrated that ICS is an alternative technique to foci counting, where the clustering involved in protein recruitment during DNA damage can be fully captured on the molecular level. ICS measures the degree of clustering and considers the low intensity signal contributions that are often lost during foci segmentation. Our findings correlate with prior studies characterizing foci formation/clustering and the DNA damage response using advanced optical techniques such laser micro-irradiation^47^ and super resolution microscopy^22^. Advantageously, ICS can be implemented on any fluorescence image as long as the square relative fluorescence intensity fluctuations are detectable above noise fluctuations. ICS does not require laser micro-irradiation to induce detectable clustering^23,47,48^. Also, it overcomes the complications introduced when using pre-extraction in immunofluorescence^49^, where soluble protein is removed through permeabilizing the cell for an arbitrary amount of time, likely removing some recruited protein as well. Thus, ICS enables simple, standard immunofluorescence labeling techniques. Other approaches that increase image resolution, such as super-resolution microscopy, will enhance the identification of single molecules and could be used to detect dimers with enough resolution enhancement. However, the definition of clustering would remain unclear as boundaries for each cluster (or foci) would have to be established, which requires subjective interpretation of the image. The overall approach, using a combination of conventional fluorescence microscopy, antibody labeling, and ICS analysis, provides a molecular level understanding for characterizing protein recruitment and signaling in a variety of applications from capturing the DNA damage response to evaluating cancer therapies. We expect adoption of this approach will both lead to a more objective measure of DNA damage and provide a tool to evaluate the role of every DDR protein during the DNA damage response.

## Data Availability

All data and analysis code are available upon contacting the corresponding author. Code for performing ICS analysis is deposited at: github.com/dubachLab/ICS.

