## Supplementary Figures for "Image Correlation Spectroscopy is a Robust Tool to Quantify Cellular DNA Damage Response"

### Supplemental Information

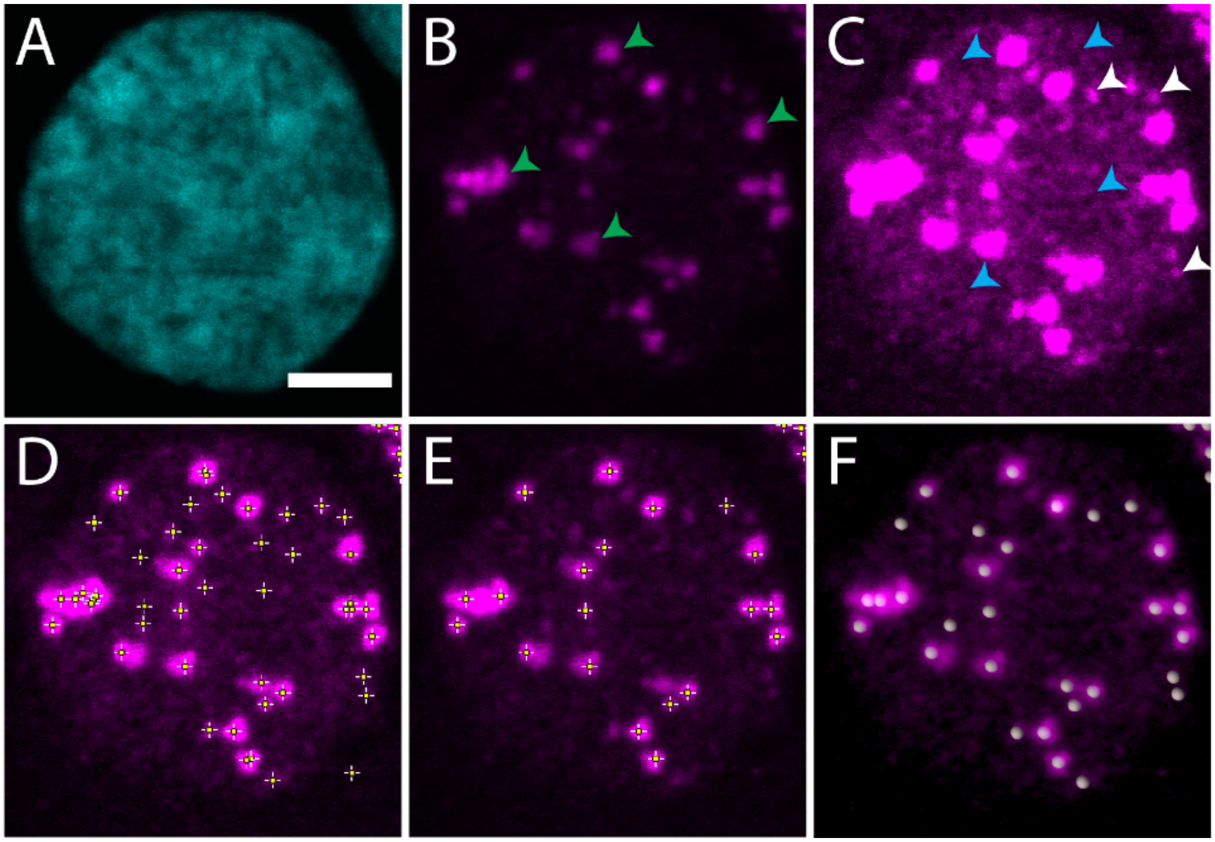


**Figure S1.** Foci Counting in a SKOV3 nucleus treated with 10 mM olaparib. (A) DAPI channel with 5 µm scale bar, (B) γH2AX channel showing larger foci (green arrows) and (C) Intensity adjusted γH2AX channel showing lower intensity foci (white arrows) and higher background intensities (blue arrows). (D-E) Foci counting with Find Maxima tool in ImageJ with a threshold of D) 600 and E) 300, (F) Foci counting using Imaris Spots with an automatic threshold.

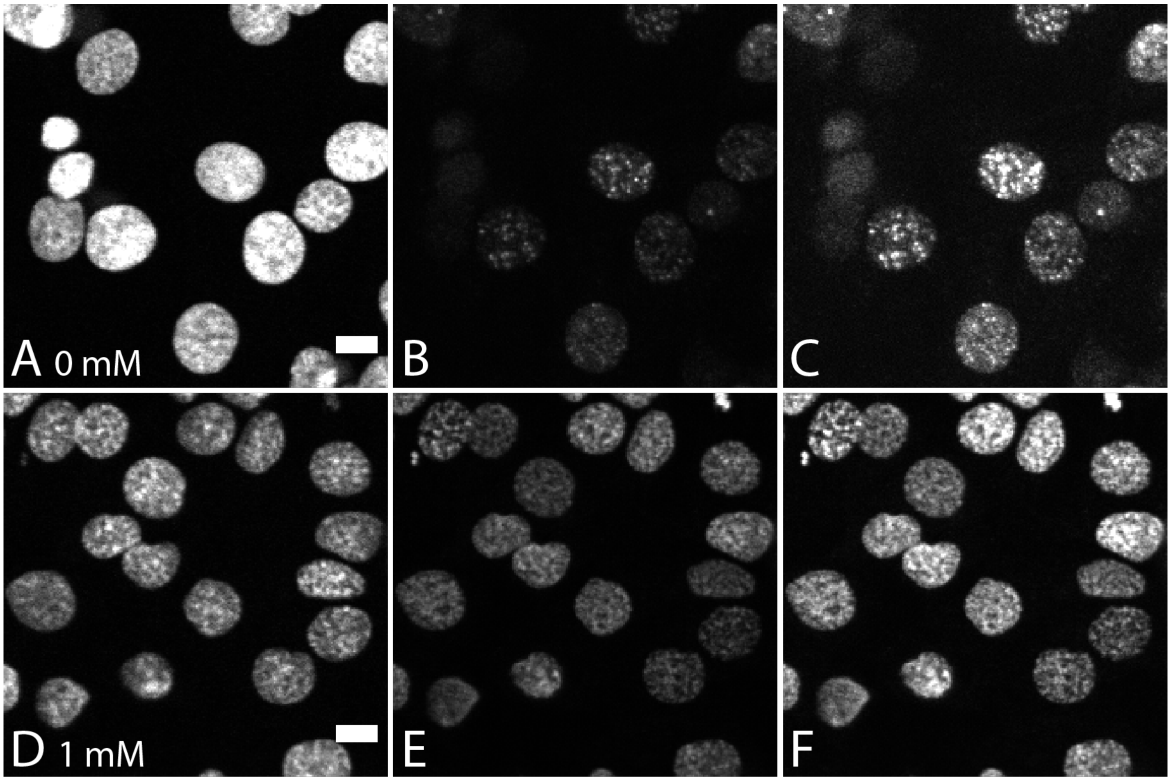


**Figure S2.** Difference between γH2AX expression in control cells (A-C) and 1 mM MMS treated cells (D-F). A) & D) DAPI channel, B) & E) γH2AX channel, C) & F) γH2AX channel with intensity adjustment. Scale bar is 10 μm.

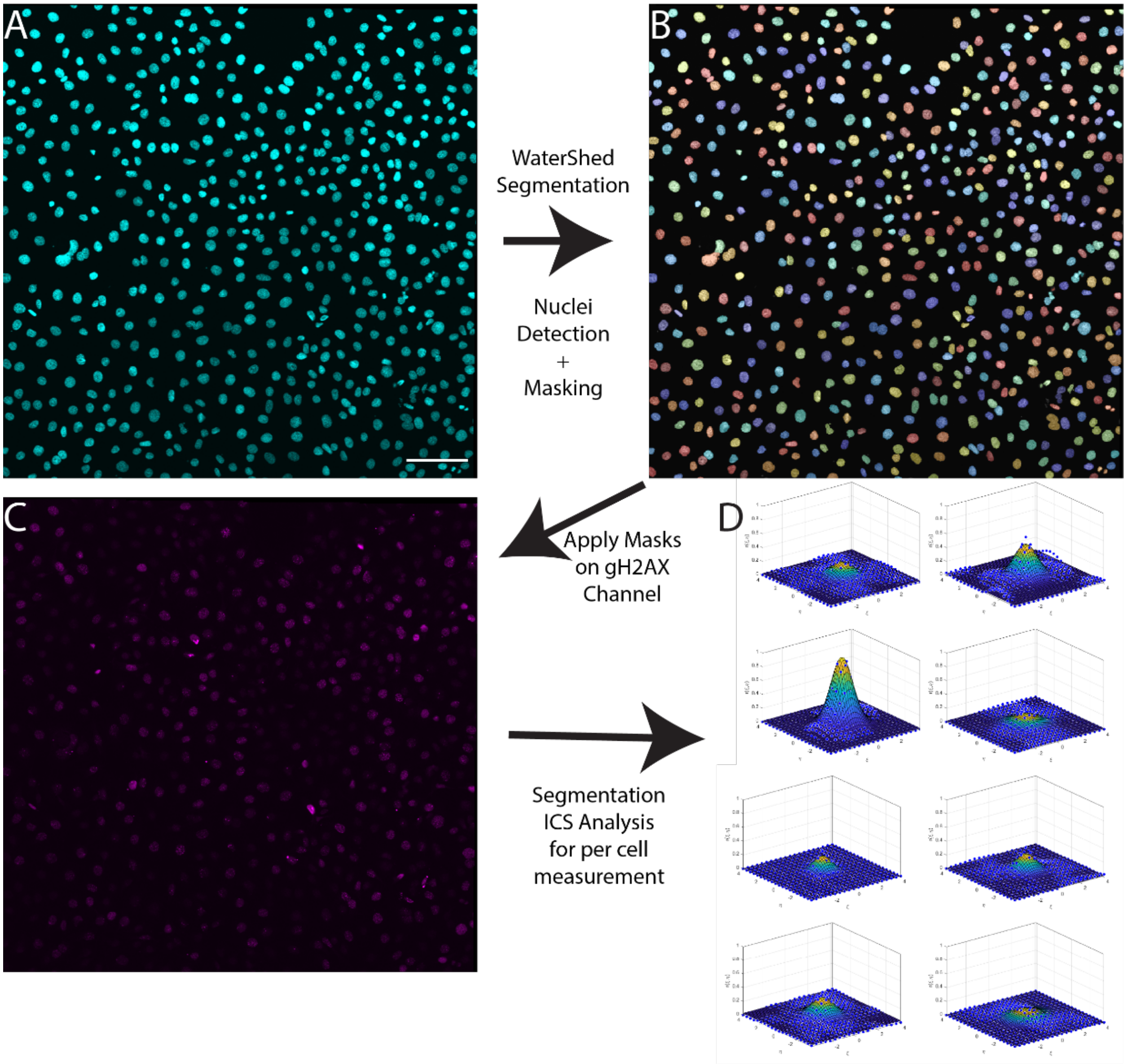


**Figure S3.** ICS Analysis of large stitched confocal microscopy images. (A) DAPI channel with 100 µm scale bar, (B) Segmentation of DAPI stained nuclei in A) using a watershed algorithm in MATLAB for nuclei detection and splitting to create nuclei masks, (C) γH2AX channel that was analyzed using the nuclei masks from B), (D) Segmentation ICS γH2AX spatial correlation functions for 8 nuclei that were displayed in C). Channels were not intensity adjusted in this image.

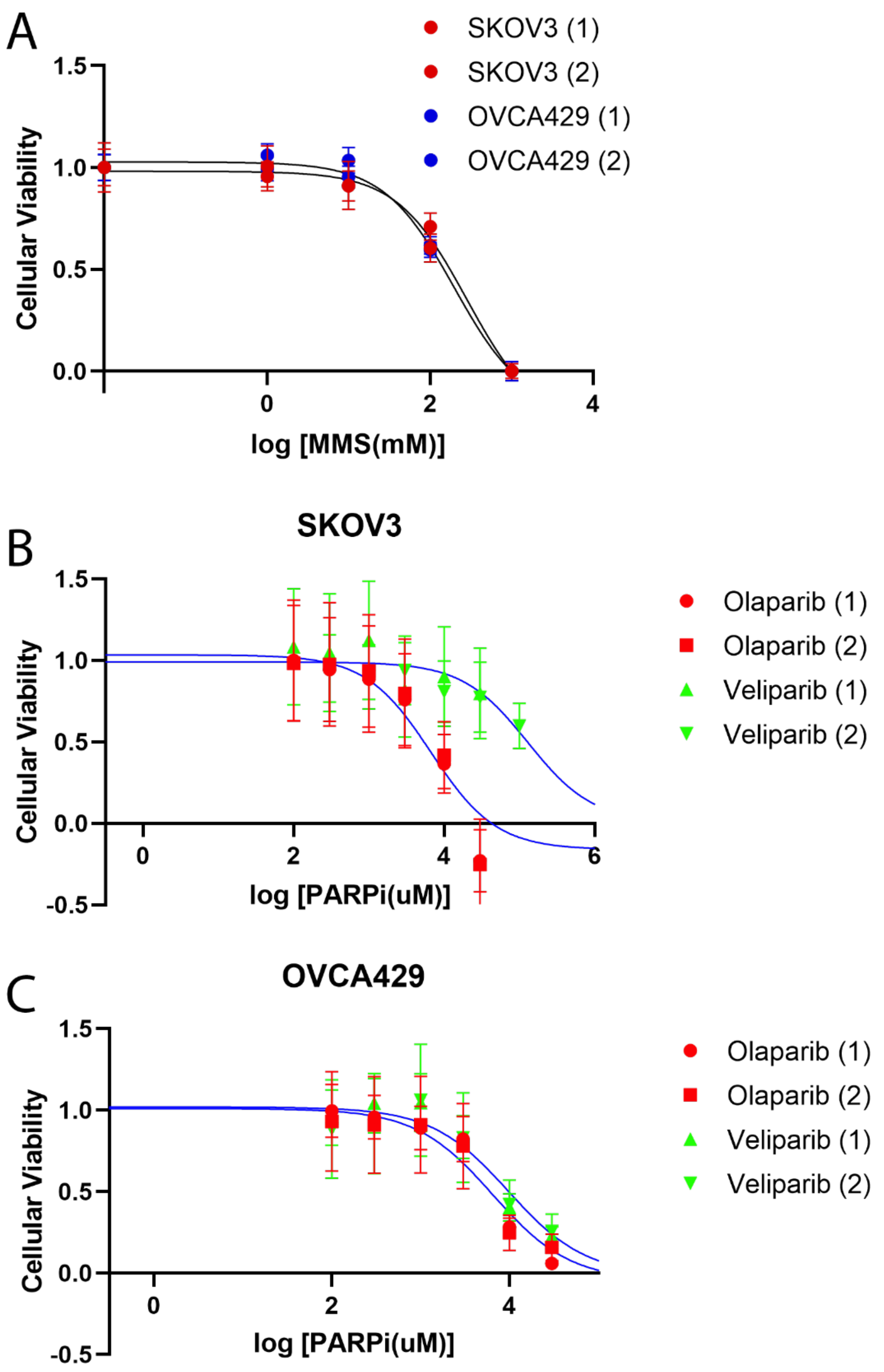


**Figure S4.** Cell dose dependence response to DDR agents. (A) SKOV3 (red markers) and OVCA429 (blue markers) dose dependent response to MMS. (B) SKOV3 dose dependent response to olaparib (red markers) and veliparib (green markers). (C) OVCA429 dose dependent response to olaparib (red markers) and veliparib (green markers). Signal was averaged over 5 replicate wells and normalized to blank (no cells) as well as to the levels of untreated wells. A sigmoidal response curve was fit to the average of two experiments. Error bars are standard deviation.

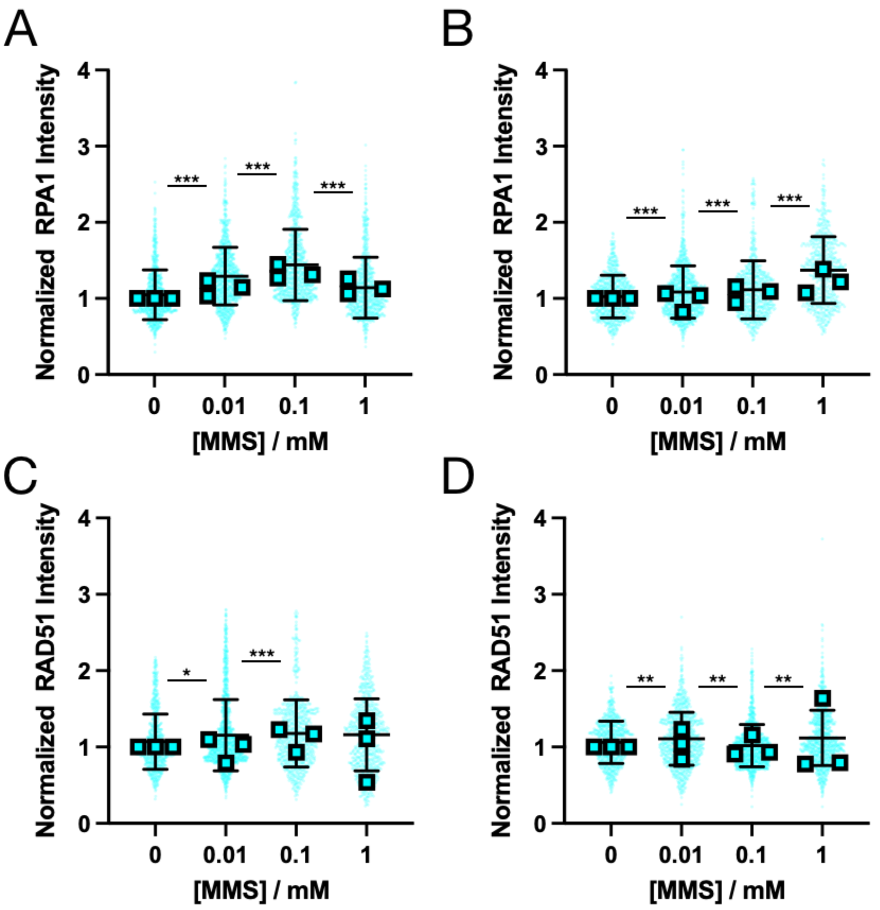


**Figure S5.** RPA1 and RAD51 intensity in SKOV3 and OVCA429 cells treated with different MMS concentrations. (A-B) Normalized RPA1 Intensity in A) SKOV3 cells and B) OVCA429 cells, (C-D) Normalized RAD51 Intensity in C) SKOV3 cells and D) OVCA429 cells across different MMS concentrations. Results are pooled from 3 independent experiments where each set of concentrations were normalized to the relevant control. N=1600-1872 for RPA1 SKOV3 and N=1320-2198 for RPA1 OVCA429. N=1597-2241 for RAD51 SKOV3 and N=1822-2311 for RAD51 OVCA429. * indicates $p<0.05$, ** indicates $p<0.001$, *** indicates $p<1X10^{-10}$ via one-way ANOVA significance testing against the control. Shown are individual cells, average for each biological repeat (n=3, squares), and average with standard deviation.

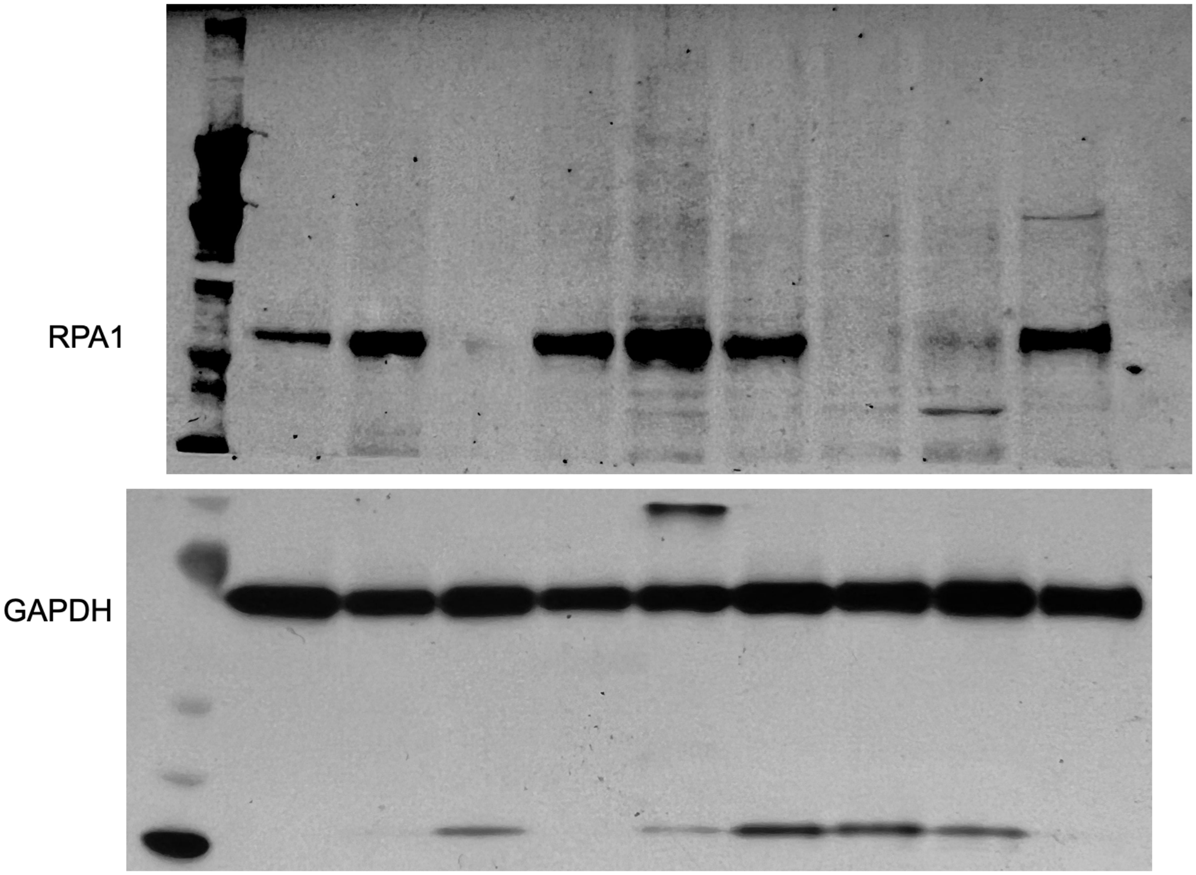


**Figure S6.** Uncropped western blots for RPA1 and GAPDH, corresponding to Figure 6.
